# Early dopamine elevation modifies the behavioral consequences of maternal separation in female and male juvenile mice

**DOI:** 10.64898/2026.10.07.757433

**Authors:** Ana Luiza de Araujo Lima Reis, Paula Bertu Valverde, Sofia Caetano Avritzer, Lorena Terene Lopes Guerra, Bruna Lopes Resende, Muiara Aparecida Moraes, Laila Blanc Arabe, Flávio Afonso Gonçalves Mourão, Márcio Flávio Dutra Moraes, Renan Pedra Souza, Bruno Rezende Souza

## Abstract

Alterations in dopaminergic signaling and exposure to early-life stress are both implicated in the developmental origins of psychiatric disorders, yet little is known about how these factors interact during sensitive periods of brain maturation. We investigated whether increased dopamine availability during early postnatal development modifies the behavioral consequences of maternal separation in female and male C57BL/6 mice. From postnatal day (PD) 1 to PD14, mice received L-Dopa plus benserazide, maternal separation (3 h/day), their combination, or the corresponding control conditions. We assessed mice behavior between PD28 and PD32 using novel object recognition, open-field, elevated plus-maze, marble-burying, and forced-swim tests, followed by hippocampal volumetric analysis. Early L-Dopa exposure produced sex- and behavior-specific effects. In females, L-Dopa increased rearing without altering horizontal locomotion and modified open-arm exploration and stress-coping behavior, whereas in males it increased open-arm exploration and reduced climbing frequency. Maternal separation produced distinct effects in females and males and reduced repetitive digging/burying behavior in both sexes. Importantly, maternal separation frequently modified rather than potentiated the effects of L-Dopa. In both sexes, combined maternal separation and L-Dopa opposed several L-Dopa-associated alterations in open-field, elevated plus-maze, and forced-swim behaviors. Recognition memory was largely preserved across experimental conditions. L-Dopa also reduced hippocampal volume selectively in females, whereas hippocampal volume was higher in females exposed to maternal separation plus L-Dopa than in those receiving L-Dopa alone. Multivariate analyses revealed overlapping rather than treatment-specific behavioral clusters, indicating substantial individual heterogeneity and sex-dependent shifts in multidimensional behavioral profiles. Together, these findings demonstrate that transient alterations in dopamine availability and maternal separation interact non-additively during early postnatal development. Their consequences depend on sex, behavioral domain, and developmental timing, supporting a model in which neurochemical and environmental perturbations dynamically shape behavioral trajectories rather than simply producing cumulative developmental damage.

## 1. Introduction

Alterations in dopaminergic signaling and exposure to stress during development are central to complementary hypotheses of vulnerability to several psychiatric disorders (Ijomone et al., 2024). In schizophrenia-spectrum conditions, elevated striatal dopamine synthesis has been detected even before the onset of psychosis (Howes et al., 2011), whereas altered striatal dopamine transporter availability has been reported in major depressive disorder (Pizzagalli et al., 2019). Childhood adversity is also associated with the subsequent onset of mood, anxiety, and substance use disorders, indicating that stressful experiences during development may contribute to vulnerability across psychiatric disorders (Green et al., 2010). Recent co-twin analyses indicate that associations between childhood adversity and adult psychiatric diagnoses persist after accounting for shared genetic and environmental factors, particularly following multiple adverse experiences or sexual abuse (Daníelsdóttir et al., 2024). Connecting these lines of evidence, a multimodal neuroimaging study linked major depression and childhood sexual abuse to region-specific abnormalities in dopamine transporter availability, with abuse history also associated with reduced frontostriatal connectivity (Borchers et al., 2026). These findings raise the possibility that early adverse experiences and dopaminergic alterations converge during brain maturation to influence psychiatric vulnerability.

Dopamine contributes directly to brain development through receptor-dependent effects on cellular processes involved in circuit formation. Experimental studies have shown that dopaminergic signaling regulates progenitor-cell proliferation in the embryonic forebrain and the development of GABAergic neurons (Crandall et al., 2007; Ohtani et al., 2003; Souza et al., 2024; Souza and Tropepe, 2011). Dopaminergic modulation also matures postnatally (Bjerke et al., 2025). For example, in rats, the effects of D2 receptor activation on prefrontal interneuron excitability differ between preadolescence and adulthood (Tseng and O’Donnell, 2006). These findings suggest that changes in dopamine availability during sensitive developmental periods may influence circuit organization and subsequent behavior (Graham et al., 2015; Kostrzewa et al., 2016, 2004; Lieberman et al., 2018; Moraes et al., 2026; Okubo et al., 2018). Consistent with this possibility, our previous work demonstrated that an early postnatal increase in dopamine produces different behavioral outcomes in juvenile and adult female and male mice (de Matos et al., 2020, 2018). Dopaminergic activity during early development may therefore shape behavioral trajectories in ways that depend on sex and developmental stage.

Early-life stress can likewise modify brain maturation and produce behavioral consequences that emerge during adolescence or persist into adulthood (Daníelsdóttir et al., 2024; Godoy et al., 2018; Hu et al., 2024; Scarpa et al., 2025; Wang et al., 2020). In humans, longitudinal neuroimaging has associated childhood maltreatment with altered developmental trajectories of the hippocampus and amygdala, linking early adversity to the maturation of regions involved in memory and emotional regulation (Whittle et al., 2013). More recently, an analysis of 7,036 participants in the Adolescent Brain Cognitive Development study associated greater childhood adversity with smaller increases in hippocampal volume during early adolescence (Breslin et al., 2024). Experimental evidence show that early-life stress in mice induces persistent epigenetic changes in hypothalamic regulation, accompanied by increased corticosterone secretion and altered memory and stress-coping behavior (Murgatroyd et al., 2009). Disrupted maternal care can also produce behavioral alterations that emerge during adolescence and become more pronounced in adulthood, with different outcomes in females and males (Goodwill et al., 2019). These findings support assessing behavior before adulthood to identify the emergence of stress-related alterations and their variation between sexes.

Stress and dopaminergic signaling interact bidirectionally. Acute stress can evoke dopamine release, and early postnatal rearing conditions can produce lasting changes in the dopaminergic system (Anderson et al., 2026; Brake et al., 2004; Pruessner, 2004). Recent mouse experiments show that early-life stress impairs dopamine signaling from the ventral tegmental area (VTA) to the basolateral amygdala and weakens the coordination of amygdala–prefrontal network activity (Stone et al., 2025). Conversely, manipulating dopaminergic activity alters stress-related behavioral responses. Optogenetic activation of specific VTA dopamine pathways can increase susceptibility to social defeat, whereas activating dopamine neurons in a chronic mild stress model can reverse selected behavioral deficits (Chaudhury et al., 2013; Tye et al., 2012). More recent experiments found that activating nucleus accumbens-projecting dopamine neurons during social defeat promoted resilience when stimulation was delivered randomly or timed to fighting back (Willmore et al., 2022). Thus, the effects of enhanced dopaminergic activity depend on the targeted circuit, stimulation pattern, and stress context. This reciprocal relationship provides a rationale for investigating whether dopamine availability during early adversity modifies its subsequent behavioral consequences.

We hypothesized that increasing dopaminergic tone during early-life stress modifies its behavioral consequences in the prepubertal period, with effects that may differ between females and males. To test this hypothesis, we examined maternal separation, L-Dopa plus benserazide administration, and their combination from postnatal day (PD) 2 to PD14 in C57BL/6 mice, using L-Dopa treatment as a pharmacological strategy to increase dopamine availability. We assessed locomotor and exploratory activity, anxiety-related behavior, coping responses to an inescapable stressor, repetitive digging behavior, and recognition memory between PD28 and PD32, and additionally examined hippocampal volume. Thus, we investigated whether pharmacological enhancement of dopaminergic signaling during early postnatal development modifies the effects of maternal separation on subsequent behavioral expression.

## 2. Materials and Methods

### 2.1 Animals

Male and female C57/BL6 mice (C57BL/6JUnib; 8–12 weeks of age) were purchased from the Animal Facility of the Universidade Federal de Minas Gerais (CEBIO, UFMG, Brazil) for mating. We raised the mice in polypropylene cages (27 × 17 × 12 cm), provided with wood shavings, housed two per cage, and maintained in a climate-controlled room with a 12 h–12 h light/dark cycle (lights on from 8 am to 8 pm), temperature at 22 ± 2 °C, and 40 –70% relative humidity. Mice had free access to food (Nuvilab CR1-Nuvital) and water.

### 2.2 Animal Manipulation and Mating

We stimulated mating by placing two females with one male. After 7 days, we removed the male from the cage. On the 19th day of gestation, we housed each female individually and daily inspected for the presence of pups. We designated the day of birth as postnatal day 1 (PD1). We standardized all postnatal handling: we kept litter size constant (4-6 pups) to avoid variation in maternal care, minimized pup handling time, and weaned the mice at PD 21. In the period between weaning and the tests, we habituated the mice to the experimenter, who performed the cleaning of the boxes (every 5 days) and gently handled them similarly to what would happen during the behavioral testing.

All animal experiments were performed following the U.S. National Institutes of Health (NIH) Guide for the Care and Use of Laboratory Animals (NIH Publication No. 80–23, revised in 1996) and were approved by the Animal Use Ethics Committee of the Federal University of Minas Gerais (CEUA 86/2016). All efforts were made to minimize the number of animals used and to prevent avoidable discomfort.

### 2.3 Experimental Design

At P2, litters were randomly assigned to their experimental groups. Female and male mice were divided into 6 groups: (1) only handled (Naïve); (2) saline-injected (Saline); (3) L-Dopa + benserazide injected (L-Dopa); (4) maternally separated (MS); (5) maternally separated and saline-injected (MS+Saline); (6) maternally separated and L-Dopa + benserazide injected (MS+L-Dopa) (Figure 1). All protocols were performed during the light phase of the cycle.

**Figure 1:**
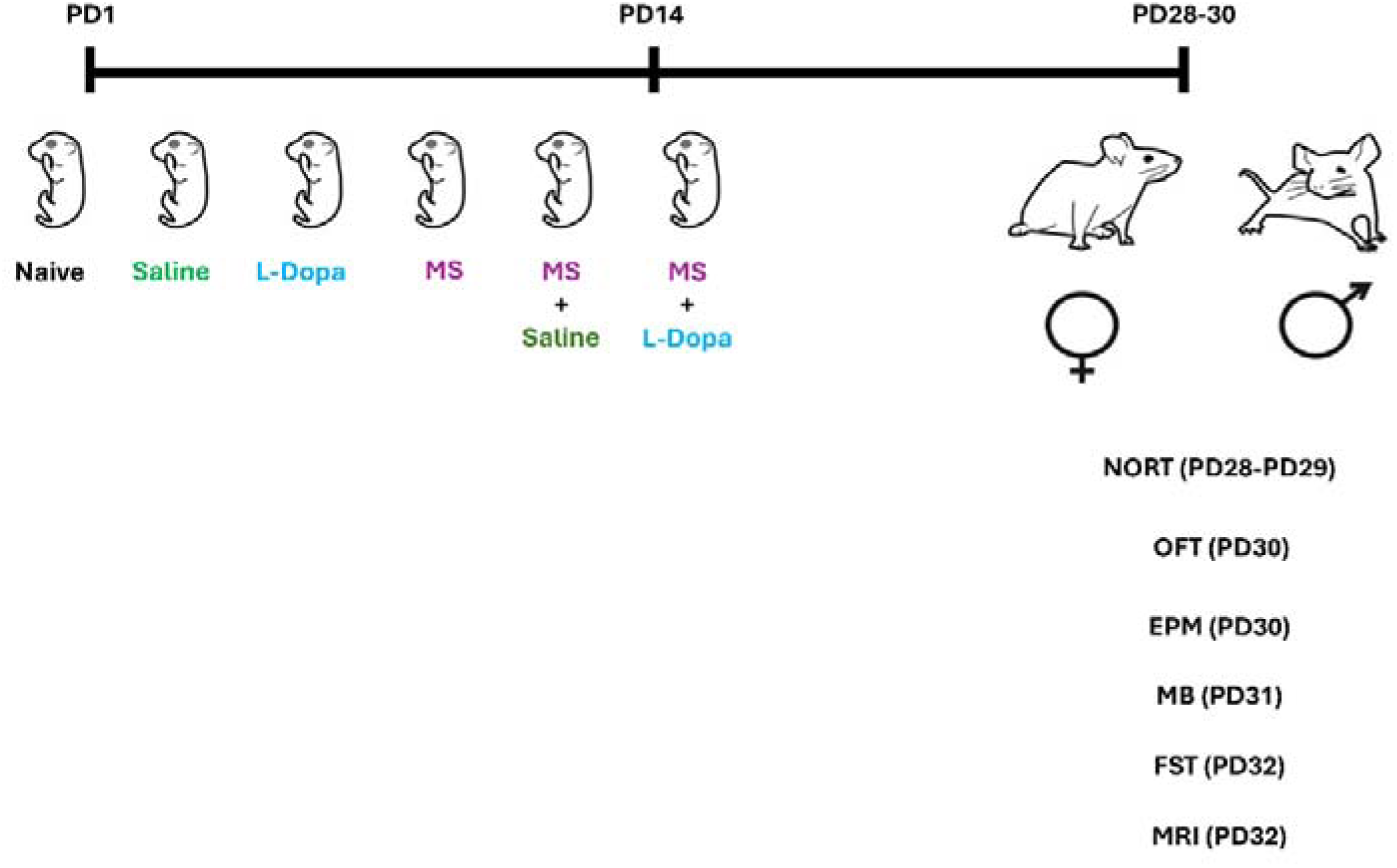
Experimental design. Male and female C57BL/6JUnib mice aged 6– 8 weeks were housed together for mating for four days. Pregnant females were housed in pairs and separated on the 19th day of pregnancy for birth. From the first day of birth (PD1) to the 14th day of age (PD14), pups were weighed and intraperitoneally injected once daily with Saline or L-Dopa, or received daily maternal separation (MS), MS+Saline, or MS+L-Dopa. Naïve, Saline, and L-Dopa manipulations lasted an average of 1–2 min, and the pups were returned to the cage with the mother immediately after the manipulation. The maternal separation protocol lasted 3 hours per day, from PD 2–14 (14:00-17:00). Pups from the same litter were removed from the home cage and placed together into cylindrical plastic containers with bedding. These containers were placed in a water bath (32°C ± 2°C) inside an acoustically isolated wood box in a different room. MS+Saline and MS-L-Dopa pups were injected daily just before the MS protocol. Dams remained in home cages, and pups were returned to their home cages after separation. The pups were raised together with their mothers until PD 22, when they were separated by sex and treatment. All mice underwent behavioral tests: novel object recognition test (NORT) (PD28-PD29), open field test (OFT) (PD30), elevated plus maze (EPM) (PD30), marble burying (MB) (PD31), and forced swim test (FST) (PD32).

### 2.4 Maternal Separation (MS) Protocol

Pups from MS, MS+Saline, and MS+L-Dopa groups were separated from their dams daily for 3 hours per day, from PD 2–14 (14:00-17:00). Pups from the same litter were removed from the home cage and placed together into cylindrical plastic containers (50mm diameter × 57 mm height) with bedding (wood shavings). These containers were placed in a water bath (32°C ± 2°C) inside an acoustically isolated wood box in a different room. Dams remained in home cages, and pups were returned to their home cages after separation.

### 2.5 Drug Administration

Pups in the Saline and MS+Saline groups were injected intraperitoneally with isotonic saline solution (0.9%) (Saline: 350 microliters/30g body weight; 29G x 1/2 needle). Saline pups were injected daily at 14:00 from PD 2–14 and returned to their home cages. MS+Saline pups were injected daily just before the MS protocol.

Pups from the L-Dopa and MS+L-Dopa groups were injected intraperitoneally with the DA precursor L-Dopa+benserazide (50/25 mg/kg; Sigma Aldrich D1507 and B7283, respectively; 350 microliters/30g body weight; 29G X 1/2 needle). We determined the L-Dopa concentration according to previous studies (including studies from our laboratory) (de Matos et al., 2018; Eskow Jaunarajs et al., 2010; Espadas et al., 2012; Garzon-Aburbeh et al., 1986; Nikolaus et al., 2016, 2014; Nikulina et al., 1991; Rodgers et al., 1994; Tronci et al., 2013; Y. et al., 2002; Zhang et al., 2003). We injected L-Dopa pups daily at 14:00 from PD2-14 and returned them to their home cages. We injected MS+L-Dopa pups daily just before the MS protocol.

### 2.6 Behavior Tests

We performed behavioral protocols in prepubescent (PD28-PD32) male and female C57/BL6 mouse offspring. The experimental design for behavioral tests was based on the puberty markers for C57/BL6 mice described in the scientific literature and on the general three-stage classification, in which days 21 to 34 of age are classified as prepubescence/early adolescence, days 34 to 46 of age as mid-adolescence, and days 46 to 59 of age as late adolescence (McCormick and Mathews, 2007; Tirelli et al., 2003). In female C57/BL6 mice, vaginal opening typically occurs around P28, while estrous cycling only begins around P35 (Iavicoli et al., 2004; Jouhanneau et al., 2015; McDermott et al., 1978; Mello et al., 2015; Safranski et al., 1993). In male C57/BL6 mice, preputial separation occurs around P25, with testosterone levels beginning to rise around P30 and peaking around P50 (BARKLEY, 1977; Jean-Faucher et al., 1985; Tyl et al., 2008).

All the experiments were performed during the light phase of the cycle, between 8:00 and 12:00 a.m. The behavioral testing room was illuminated at 365-lux. After each test, we cleaned the equipment with 70% ethanol to eliminate olfactory cues. To habituate the mice to behavioral testing room conditions, we placed mouse cages in the laboratory for 30 minutes before the experiment. We handled the mice by grabbing their tails, placing them on the experimenter’s palm, and placing them in the testing apparatus by grabbing their tails (total handling time ∼15-20 s).

All animals underwent all behavioral tests, from PD28 to PD32, in the following order: new object recognition test (PD28-PD29), open field test (PD30), elevated plus maze (PD30), marble burying (PD31), and forced swim test (PD32).

### 2.7 Novel Object Recognition Task (NORT)

The protocol used was described by Gresack and Frick, 2006. On the first day of the protocol, mice were habituated to the training arena (50cm x 40cm x 20cm plastic box), where they freely explored the space without stimulus objects for 10 minutes (habituation session). Twenty-four hours (24 h) after habituation, mice were trained in a novel object-recognition task (training session). Mice were positioned for 1 minute in the box for visual exploration of two identical objects. We measured the time spent investigating both objects until total exploration reached 30 s. We quantified exploration by every time the animal touched the object with its front paws, nose, or vibrissae. In the first test (short-term memory), we replaced one of the objects familiar to the animal with a new object one and a half hours (1.5 h) after training. In the second test (long-term memory), the animals were exposed to a familiar object and a new object twenty-four hours (24 h) after training.

The box was sanitized with 70% alcohol at each new test to prevent olfactory adaptations. For object selection, we considered appropriate weight and height to avoid both animal movement on the object and movement of the object from its initial position.

The discrimination index (*d2*) of the new object was calculated by dividing the exploration time of the new object (*d1*) by the total exploration time (*e2*). (*d2 = d1/e2*)

### 2.8 Open Field Test (OFT)

The OFT assesses spontaneous locomotor activity in a new environment (Kulesskaya and Voikar, 2014; Seibenhener and Wooten, 2015), and decreased time in the center of the open field can be used to assess anxiety-like behavior (Gwendolyn G. Calhoon and Tye, 2015; Prut and Belzung, 2003). The apparatus (LE 8811 IR Motor Activity Monitors PANLAB, Harvard Apparatus; Spain) consists of a 22 x 22 x 30 cm acrylic box with transparent walls and a black floor, coupled to an external infrared system. At the beginning of the test, we placed the animals individually in the center of the arena and measured locomotor and exploratory activity for 20 minutes. We used the automated system Actitrack v2.7.13 to evaluate the total distance traveled horizontally by the animal, the exploration time in the arena divisions (center and periphery), as well as the number of rearings – a parameter used for vertical exploration (Spielewoy et al., 2000).

### 2.9 Elevated Plus Maze (EPM)

The EPM is a test that induces an approach/avoidance conflict to assess anxiety-like responses in rodents (Walf and Frye, 2007). The apparatus consists of two open arms (30 × 6 cm) and two closed arms (30 × 6 × 15 cm) crossed in the middle, perpendicularly to each other, and elevated 40 cm above the floor. We placed the mice in the central square, facing one of the open arms, and allowed them to move freely for 5 min. We used the number of entries into the open arms and the time spent in the open arms as indices of anxiety-like behavior (Pellow et al., 1985). We also analyzed ethological parameters such as rearing, stretching (forward elongation of head and shoulders followed by retraction to the original position), and head-dipping (lowering the head by the sides of the open arms to the floor) to explore the complexity of mouse behavior and anxiety-like behavior (Bailey and Crawley, 2009; Espejo, 1997; Rodgers et al., 1997; Rodgers and Johnson, 1995). Rearings and stretchings were counted regardless of where they occurred, while head-dipping is possible only when the animal is on one of the open arms.

### 2.10 Marble Burying Test (MB)

The marble burying test is commonly used to assess repetitive behaviors/compulsive-like behaviors (Angoa-Pérez et al., 2013). Animals were individually placed in an acrylic box (37cm x 21cm x 14cm) containing 20 standard marbles (assorted styles and colors, 15 mm diameter, 5.2 g each), separated from one another. The marbles were placed on top of the shaving (depth 5cm) in 5 rows of 4 marbles. The duration of the test was 20 min, and we counted the number of balls buried in the shaving, at least 2/3, at the end of the test (Thomas et al., 2009).

### 2.11 Forced Swim Test (FST)

We used the Forced Swim Test to evaluate despair/depressive-like behavior in rodents, as reviewed by Can et al. (2012), with minor modifications (Can et al., 2011a; Porsolt et al., 1977). Mice were individually transferred to a vertical glass cylinder (17 cm inner diameter × 27 cm height) filled with water and were forced to swim for 6 minutes (Can et al., 2011b). We monitored the water temperature throughout the test and maintained it between 25 and 27 °C. We analyzed the 6-minute test, evaluating latency to the first episode of immobility (when the animal moves as little as possible, just to stay on the surface), total immobility time, and climbing (vertical movements of the forepaws directed against the wall of the cylinder). Reduced immobility latency, increased total immobility time, and reduced climbing and swimming behaviors indicate despair/depressive-like behavior in animals (Petit-Demouliere et al., 2005).

### 2.12 Hippocampal Structural Magnetic Resonance Imaging

Following all behavioral tests, mice were anesthetized with isoflurane (3% induction and 1.5% maintenance) and oxygen (1.5 l/min) administered via a mask. We gently secured the mice’s heads and bodies within the MRI coil to minimize sudden movements during image acquisition. Imaging was performed using a 4.7T NMR magnet (Oxford Systems) controlled by a UNITY Inova-200 imaging console (Varian). The scanning protocol consisted of T2-weighted (TR=300ms, TE=50ms) spin-echo multislice scans, generating 20 consecutive coronal slices, each 1 mm thick.

Image acquisition utilized a resolution of 256 × 128 voxels within a slice measuring 3.2 × 3.2 cm. We then assessed images using manually constructed masks for the entire hippocampus and quantified volumes using the NIH ImageJ program (https://imagej.nih.gov/ij/).

### 2.13 Statistical analysis

We performed statistical analyses and created graphs using SigmaPlot 13.0 and GraphPad Prism 10.2.3, respectively. We assessed the main effects (treatment and MS) using Two-Way ANOVA (TW ANOVA) with Tukey’s test for multiple comparisons. Significance was set at p < 0.05 for all analyses.

#### 2.13.1 Multivariate behavioral analyses

Multivariate analyses were used to characterize behavioral patterns that may be unobservable through conventional comparison of group means. We aimed to identify sex- and treatment-specific relations within each behavioral test and to assess whether animals could be grouped into distinct multidimensional behavioral profiles. We performed all analyses separately by sex, and standardized variables as z-scores before correlation analysis, non-hierarchical clustering, UMAP visualization, and individual-level heatmap generation. We conducted all analyses in R (version 4.5.2) and RStudio (2026.01.0 Build 392) using the base “stats” implementation of k-means clustering, “uwot” for UMAP dimensionality reduction, and “ComplexHeatmap” and “ggplot2” for data visualization.

#### 2.13.2 Within-test correlation analysis

We performed correlation analysis among variables within each behavioral test, separately for sex and experimental group. Pairwise correlations were calculated using Spearman correlation tests, and p-values were adjusted using the Benjamin-Hochberg false-discovery rate. We presented Spearman’s correlation coefficient (rs) values and adjusted p-values in correlation heatmaps for all analyses.

#### 2.13.3 Non-hierarchical clustering and UMAP visualization

We evaluated behavioral organization patterns separately in males and females for each behavioral test. We applied k-means clustering to the complete standardized behavioral feature space using Euclidean distance, with the number of clusters fixed a priori at 4 (k = 4) to match the number of experimental groups. To reduce sensitivity to initial centroid placement, we ran the algorithm with 100 random initializations and kept the solution with the lowest total within-cluster variation. We used uniform manifold approximation and projection (UMAP) to visualize the multidimensional behavioral data. We calculated UMAP from the same standardized variables used for clustering, using Euclidean distance, 15 nearest neighbors, a minimum distance of 0.15, and a spread parameter of 1. We obtained cluster assignments from k-means clustering in the full multidimensional feature space, not from the two-dimensional UMAP coordinates; UMAP was used exclusively for visualization.

#### 2.13.4 Cluster behavioral profiles

To characterize the behavioral phenotype associated with each k-means cluster, we examined cluster profiles separately for each sex and behavioral test. We calculated the mean standardized value across all animals assigned to a given cluster for each variable, and present profiles as mean z-scores and standard error of the mean (SEM). Positive values indicate behavioral measurements above the sex-specific mean for that test, while negative values indicate measurements below the mean.

#### 2.13.5 Individual behavioral heatmaps

We visualized individual behavioral patterns as heatmaps, separately for males and females, within each test. Heatmap values correspond to z-scores, with the color scale centered at zero. We divided animals into four blocks based on their k-means cluster assignments.

## 3. Results

### 3.1 The increase in dopamine levels in the first 14 postnatal days induces exploratory activity only in prepubertal females, yet this effect is attenuated when combined with maternal separation

Hyperactivity of the dopaminergic system is correlated with alterations in locomotor activity. Our recent research showed that increased dopamine levels during the first 5 days of life affect locomotor/exploratory activity in both juvenile and young adult female mice (de Matos et al., 2020, 2018). In this study, we aimed to explore not only the impact of increased dopamine levels during the first 14 days of life on prepubertal mouse behavior but also potential interactions between maternal separation-induced stress and dopamine increases during this critical period. We evaluated locomotor/exploratory activity using the OFT (Fig. 1). We did not see changes in total distance or time spent in the center in prepubertal females (Fig. 2A-B). However, we observed a significant increase of 30% in the number of rearings in L-Dopa females compared to Naïve (F (2,82) = 5.396, p = 0.006; Power = 0.744) and 32% compared to Saline prepubertal females (F (2,82) = 5.396, p = 0.021; Power = 0.744) (Fig. 2C). Interestingly, there is a decrease of 19% in the number of rearings in MS+L-Dopa compared to L-Dopa prepubertal females (F (2,82) = 5.396, p = 0.013; Power = 0.744) (Fig. 2C). However, we observed no alterations in the locomotor/exploratory activity in prepubertal males (Fig. 2D-F).

**Figure 2:**
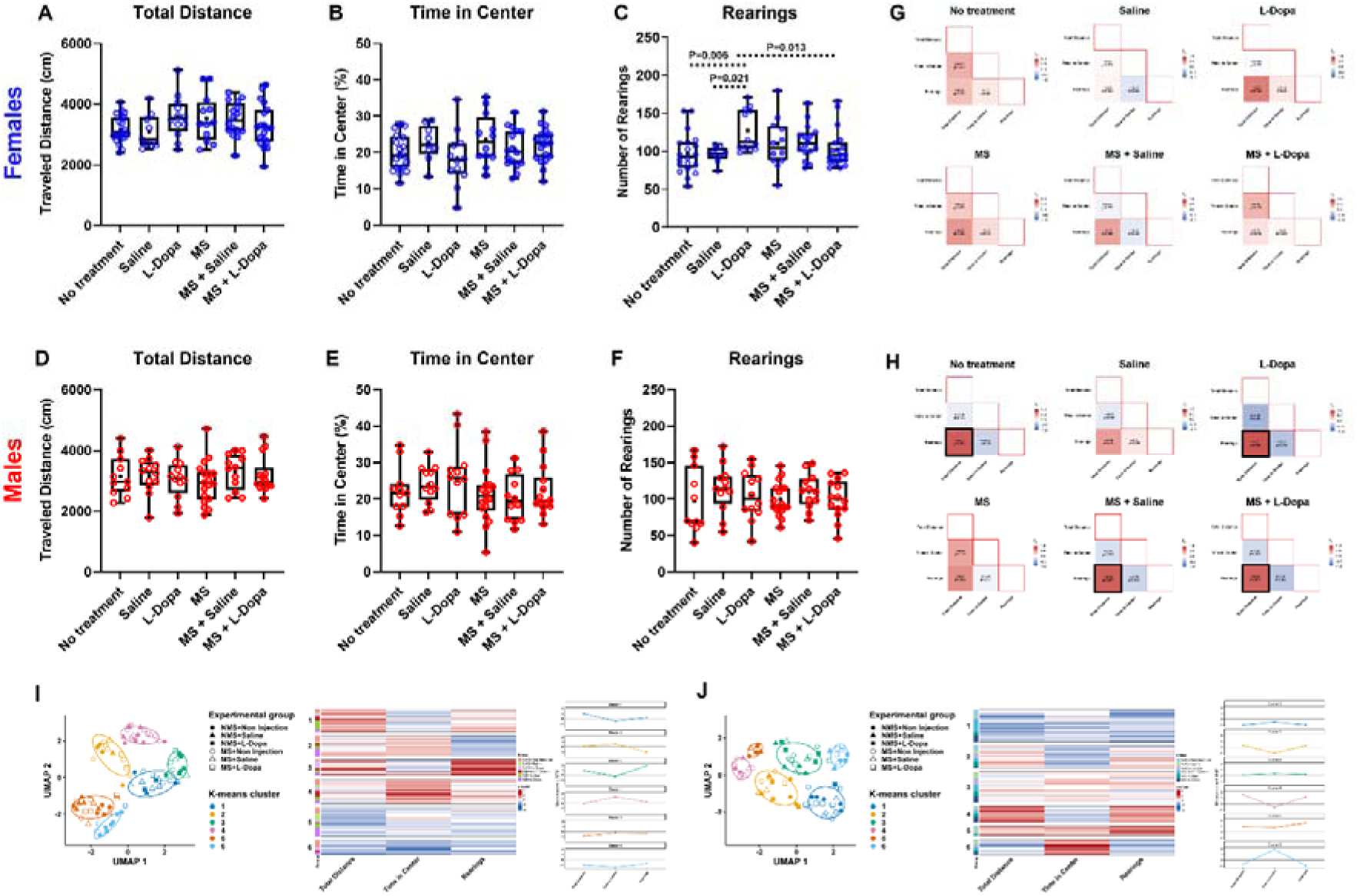
The increase in dopamine levels in the first 14 postnatal days induces exploratory activity only in prepubertal females, yet this effect is attenuated when combined with maternal separation. Exploratory and motor behavior of prepubertal C57/BL6 mice in the Open Field Test. (A-B) Prepubertal females from all experimental groups showed a similar traveled distance and time in the center. (C) Prepubertal L-Dopa females showed more rearings than Saline and Naïve females. MS+L-Dopa female mice showed fewer rearings than prepubertal L-Dopa females. (D-F) Prepubertal males from all experimental groups showed a similar traveled distance, time in the center, and number of rearings. MS: Maternal Separation. Blue circles: female mice. Red circles: male mice. Empty circles: no maternal separation. Filled circles: maternally separated. (+) Mean; (Line) Median. The main effects (L-Dopa administration and MS) were assessed with Two-Way ANOVA with Tukey’s test for multiple comparison procedures. Significance was set at p < 0.05 for all analyses. (C) Correlations between latency and immobility time in Saline, L-Dopa, SKF, and Quinpirole females and males. Color intensity indicates the degree of correlation (red indicates a positive correlation and blue a negative correlation). The heatmap shows Spearman correlations and p-values. Dark borders indicate p ≤ 0.05. (D) UMAP visualization of standardized behavioral features, analyzed separately in females and males. Colors represent the four k-means clusters, whereas symbol shapes indicate treatment groups. (E) Individual heatmaps showing standardized behavioral values within each cluster. Rows represent individual animals, columns represent TST variables, and colors indicate z-scores, with red and blue denoting values above and below the mean, respectively. Side annotations identify treatment groups, and the adjacent plots summarize the average behavioral profile of each cluster.

To evaluate the relationships among the OFT variables, we measured their correlations. There were no correlations between OFT variables in females (Fig. 2G). In contrast, untreated prepubertal males showed a correlation between total distance and number of rearings (rs = 0.75, p = 0.022). The same correlation was observed in L-Dopa (rs = 0.78, p = 0.008), MS+Saline (rs = 0.74, p = 0.017), and MS+L-Dopa (rs = 0.72, p = 0.011). (Fig. 2H)

To complement the univariate and correlation analyses, we examined the multivariate organization of OFT behavior separately in females and males. K-means clustering of the complete standardized behavioral feature space partitioned the mice into six clusters, with the number of clusters fixed a priori at (k=6). We then overlaid cluster assignments on two-dimensional UMAP projections for visualization (Fig. 2I-J). The projections showed partially distinct but overlapping regions occupied by the six clusters. Mice from the Naïve, Saline, L-Dopa, MS, MS+Saline, and MS+L-Dopa groups were intermingled throughout the projections and distributed among multiple clusters, revealing no evident treatment-specific segregation.

Individual heatmaps showed that no behavioral profile was exclusive to a group. All clusters included females from the Naïve, MS, and MS+L-Dopa groups. Females from Naïve and MS+L-Dopa were prominent in clusters 2 (intermediate distance traveled, longer time in center, and fewer rearings) and 3 (high distance traveled, intermediate time in center, and high number of rearings), while MS females were prominent in cluster 4 (intermediate distance traveled, longer time in center, and intermediate number of rearings). Interestingly, no L-Dopa females were in cluster 2. (Fig. 2I). All male groups were represented in clusters 1, 2, 4, and 6. However, males from the Naïve group were not represented in cluster 3 (intermediate distance traveled, time in center, and number of rearings), and males from MS and MS+L-Dopa were not represented in cluster 5 (higher distance traveled, time in center, and number of rearings) (Fig. 2J).

### 3.2 Maternal separation stress attenuates the anxiolytic-like effects of increased dopamine in the first 14 postnatal days

Female prepubertal mice injected with L-Dopa in the first 14 postnatal days showed an increase in the number of rearings in OFT (Fig. 2C). Dopaminergic signaling is also involved in anxiety-like behavior, and alterations in the number of rearings can indicate this behavior in mice (Gwendolyn G Calhoon and Tye, 2015; Sturman et al., 2018). Because of this, we tested this anxiety-like behavior using the EPM (Fig. 1). We observed a main effect of treatment on the number of entries in open arms in females, indicating that across groups, Saline factor decreased the number of entries in the open arms (F (2,88) = 3.689, p = 0.022; Power = 0.520). Between groups, Saline and L-Dopa prepubertal females showed an increase of 32% (F (2,88) = 2.253, p = 0.006; Power = 0.448) and 24% (F (2,88) = 3.689, p = 0.031; Power = 0.448) respectively in entries in open arms compared to Naïve females (Fig. 3A). We also observed an increase of 41% in time spent in open arms in both Saline (F (2,88) = 6.193, p = 0.042; Power = 0.817) and L-Dopa (F (2,88) = 6.193, p = 0.024; Power = 0.817) compared to Naïve females (Fig. 3B). Interestingly, MS females showed an increase of 39% in time spent in open arms compared to Naïve females (F (2,88) = 6.193, p = 0.009; Power = 0.817) (Fig. 3B). In addition, MS+L-Dopa females had a decrease of 24% in time spent in open arms compared to L-Dopa (F (2,88) = 6.193, p = 0.031; Power = 0.817) (Fig. 3B). Females showed no changes in time spent in center and in the number of head dippings (Fig. 3C and E). However, L-Dopa had a main effect on the number of rearings (F (2,88) = 6.761, p = 0.001; Power = 0.857). The number of rearings decreased by 30% in L-Dopa mice compared with MS (F (2,88) = 0.408, p = 0.014; Power = 0.05). MS+L-Dopa females showed a 32% decrease in the number of rearings compared with MS females (F (2,88) = 6.193, p = 0.014; Power = 0.05) (Fig. 3D).

**Figure 3:**
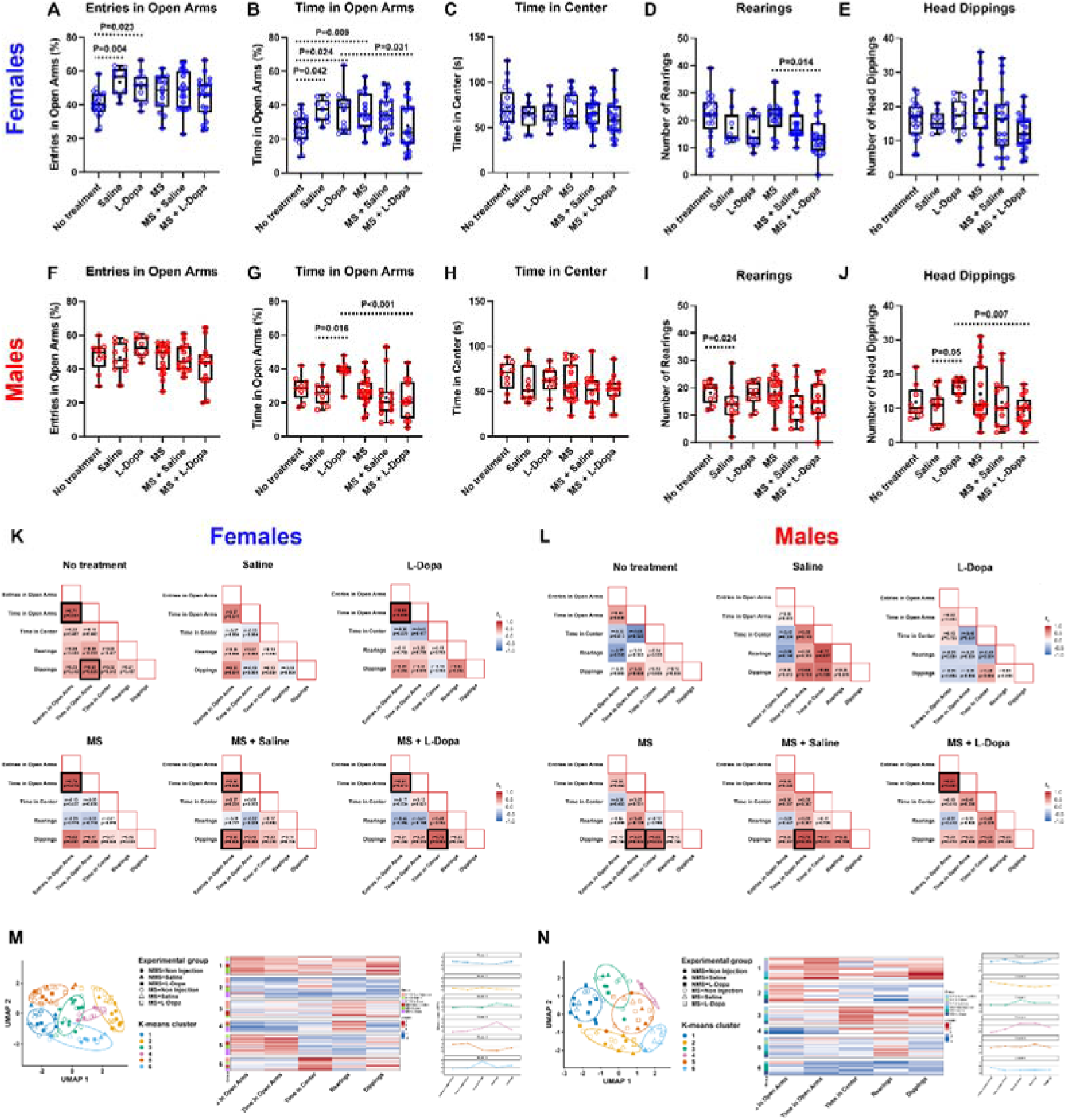
Maternal separation stress attenuates the anxiolytic-like effects of increased dopamine in the first 14 postnatal days. Anxiety-like behavior of prepubertal C57/BL6 mice in the Elevated Plus Maze test. (A) Saline and L-Dopa females showed more entries in open arms than Naïve prepubescent females. (B) Saline, L-Dopa, and MS females showed increased time in open arms compared with Naïve prepubescent females. MS+L-Dopa showed decreased time in open arms compared with L-Dopa prepubescent females. (D) The number of rearings is diminished in MS+L-Dopa compared to MS prepubescent females. (C and E) Prepubertal females in all experimental groups showed similar time spent in the center and number of head dippings. (G) L-Dopa males showed increased time in open arms compared with saline prepubescent males. MS+L-Dopa males showed decreased time in open arms compared with L-Dopa prepubescent males. (I) There is a decrease in the number of rearings in Saline males compared to Naïve prepubescent males. (J) L-Dopa males showed an increase in head dippings compared with Saline prepubescent males. MS+L-Dopa males showed fewer head dips than L-Dopa prepubescent males. (F and H) Prepubertal males in all experimental groups showed similar numbers of entries in open arms and time in the center. MS: Maternal Separation. Blue circles: female mice. Red circles: male mice. Empty circles: no maternal separation. Filled circles: maternally separated. (+) Mean; (Line) Median. The assessment of the main effects (L-Dopa administration and MS) was performed with Two-Way ANOVA with Tukey’s test for multiple comparison procedures. Significance was set at p < 0.05 for all analyses. (C) Correlations between latency and immobility time in Saline, L-Dopa, SKF, and Quinpirole females and males. Color intensity indicates the degree of correlation (red indicates a positive correlation and blue a negative correlation). The heatmap shows Spearman correlations and p-values. Dark borders indicate p ≤ 0.05. (D) UMAP visualization of standardized behavioral features, analyzed separately in females and males. Colors represent the four k-means clusters, whereas symbol shapes indicate treatment groups. (E) Individual heatmaps showing standardized behavioral values within each cluster. Rows represent individual animals, columns represent TST variables, and colors indicate z-scores, with red and blue denoting values above and below the mean, respectively. Side annotations identify treatment groups, and adjacent plots summarize each cluster’s average behavioral profile.

On the other hand, prepubescent males did not show alterations in the number of entries in open arms (Fig. 3F). There was a main effect of maternal separation in the time spent in the open arms, with a decrease of 24% (F (2,68) = 10.901, p = 0.002; Power = 0.873). We also observed a 44% increase in time spent in open arms in L-Dopa compared to Saline males (F (2,68) = 4.265, p = 0.016; Power = 0.603) (Fig. 3G). Interestingly, MS+L-Dopa had a decrease of 44% in time spent in open arms compared to L-Dopa prepubescent males (F (2,68) = 4.265, p < 0.001; Power = 0.603). Similar to females, no prepubescent males showed changes in time spent in the center (Fig. 3H), but Saline males had a decrease of 24% in the number of rearings compared to Naïve prepubescent males (F (2,68) = 4.265, p = 0.024; Power = 0.050) (Fig 3I). Saline injection had a main effect on the number of rearings, decreasing it by 26% (F (2,68) = 3.637, p = 0.002; Power = 0.508). In addition, L-Dopa males showed an increase of 62% in the number of head dippings compared to Saline (F (2,68) = 4.265, p = 0.05; Power = 0.611), but MS+L-Dopa had a decrease of 43% in the number of head dippings compared to L-Dopa prepubescent males (F (2,68) = 4.265, p = 0.007; Power = 0.611) (Fig. 3J).

We calculated pairwise Spearman correlation coefficients to assess the relationships among EPM variables (Fig. 3K,L). In females, time spent in open arms was correlated with the number of entries in open arms in Naïve (rs = 0.71, p = 0.004), L-Dopa (rs = 0.85, p = 0.005), MS (rs = 0.76, p = 0.015), MS+Saline (rs = 0.60, p = 0.026), and MS+L-Dopa (rs = 0.64, p = 0.015) (Fig. 3K). In addition, Naïve females showed a correlation between time in open arms and the number of head dippings (rs = 0.60, p = 0.026); MS+Saline showed a correlation between the number of entries in open arms and the number of head dippings (rs = 0.60, p = 0.026); MS+L-Dopa showed a correlation between time in center and the number of head dippings (rs = 0.72, p = 0.005) (Fig. 3K). Regarding males, time in open arms was correlated with the number of head dippings in MS (rs = 0.61, p = 0.035) and MS+Saline (rs = 0.79, p = 0.015) (Fig. 3L). Also, MS males showed a correlation between time in center and the number of head dippings (rs = 0.65, p = 0.035), and MS +L-Dopa showed a correlation between time in open arms and the number of entries in open arms (rs = 0.84, p = 0.004) (Fig. 3L).

UMAP visualization revealed six partially separated behavioral profiles, with treatment groups distributed across clusters (Fig. 3M,N). Individual heatmaps showed that no behavioral profile was exclusive to a group. However, females from the Naïve group were not represented in cluster 5 (more entries and time in the open arms, and less time in the center and fewer rearings). L-Dopa females were absent in cluster 4 (fewer entries and less time in the open arms, and more rearings), and MS+L-Dopa females were absent in cluster 3 (more rearings). Regarding males, the Naïve group was absent in Cluster 6 (lower levels in all parameters). L-Dopa males were prevalent in Cluster 1 (higher number of entries, time in the open arms, and head dippings) and were absent in Cluster 4 (lower number of entries, time in the open arms, and head dippings), 5 (intermediate levels in all parameters), and 6 (lower levels in the time in the open arms, time in the center, number of rearings, and head dippings). In contrast, no males from MS+L-Dopa were in Cluster 1 (Fig. 3M,N).

### 3.3 The interplay between increased dopamine levels during infancy and maternal separation stress on despair/depressive-like behavior in prepubertal males

Numerous studies have highlighted the role of dopamine in depressive behavior, making dopaminergic signaling one of the main targets for antidepressants (Otte et al., 2016; Tye et al., 2012). Bearing this in mind, we used the FST to investigate despair/depressive-like behavior in prepubescent mice (Fig. 1). There were no changes in latency immobility in prepubescent females (Fig. 4A). However, we observed a reduction in the number of immobility episodes of 41% and 45% in L-Dopa females compared to Naïve (F (2,83) = 3.292, p = 0.024; Power = 0.453) and Saline females (F (2,83) = 3.292, p = 0.017; Power = 0.453) respectively. In contrast, MS+L-Dopa females had a 77% increase in the number of immobile episodes compared with L-Dopa (F (2,83) = 3.292, p = 0.004; Power = 0.453) (Fig. 4B). L-Dopa females also showed a 64% decrease in immobility time compared to Saline (F (2,83) = 4.743, p = 0.017; Power = 0.670). MS+L-Dopa had an increase of 127% in immobility time compared to L-Dopa (F (2,83) = 4.743, p = 0.020; Power = 0.670) (Fig. 4C). There were main effects of maternal separation in the number of climbings and time of climbing, with an increase of 71% of the number of climbings episodes (F (2,83) = 23.708, p < 0.001; Power = 0.997) and of 130% in time of climbing (F (2,83) = 11.341, p = 0.001; Power = 0.888). We observed a 47% decrease in climbing episodes in Saline compared with Naïve females (F (2,83) = 2.975, p = 0.042; Power = 0.398). On the other hand, we observed an increase of 139% of climbing episodes in MS+Saline compared to Saline (F (2,83) = 2.975, p < 0.001; Power = 0.398) and of 100% in MS+L-Dopa compared to L-Dopa (F (2,83) = 2.975, p < 0.001; Power = 0.398) (Fig. 4D). In addition, MS+Saline showed an increase of 266% in climbing time compared to Saline females (F (2,83) = 0.536, p = 0.009; Power = 0.050) (Fig. 4E).

**Figure 4:**
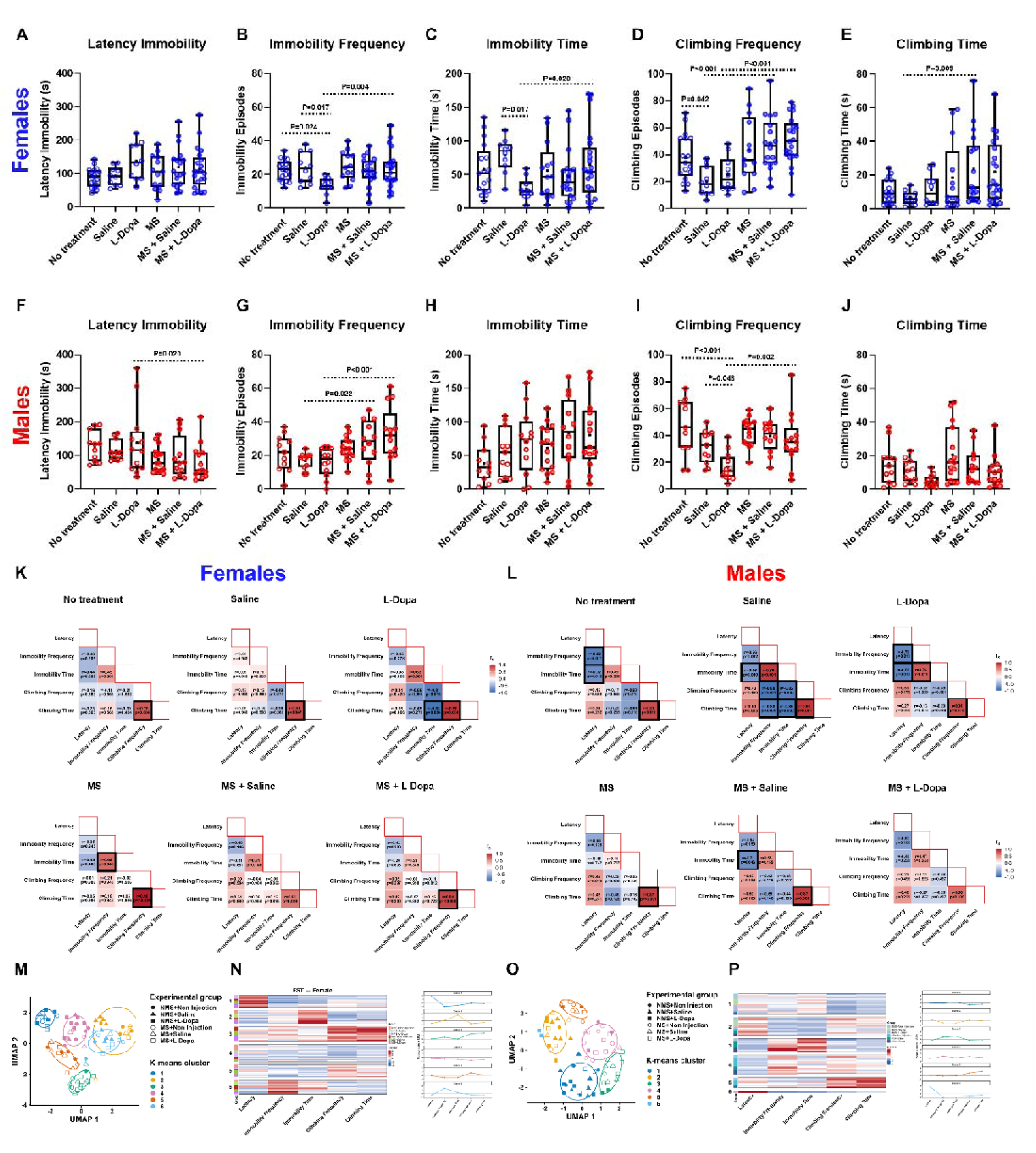
The interplay between increased dopamine levels during infancy and maternal separation stress on despair/depressive-like behavior in prepubertal males. Depressive/despair-like behavior of prepubertal C57/BL6 mice in the Forced Swim Test. (A) Prepubertal females from all experimental groups showed similar immobility latency. (B) L-Dopa females showed decreased immobility frequency compared with Naïve, Saline, and MS+L-Dopa prepubescent females. (C) L-Dopa females showed decreased immobility time compared with Saline and MS+L-Dopa prepubescent females. (D) Saline females had decreased climbing frequency compared with Naïve and MS+Saline prepubescent females. MS+L-Dopa females showed increased climbing frequency compared with L-Dopa prepubescent females. (E) MS+Saline females had increased climbing time compared with MS+Saline prepubescent females. (F) MS+L-Dopa males showed decreased immobility latency compared with L-Dopa prepubescent males. (G) MS+Saline and MS+L-Dopa males showed increased immobility frequency compared with Saline and L-Dopa prepubescent males, respectively. (I) L-Dopa males showed decreased climbing frequency compared with Naïve, Saline, and MS+L-Dopa prepubescent males. (H,J) Prepubertal males from all experimental groups showed similar immobility and climbing time. MS: Maternal Separation. Blue circles: female mice. Red circles: male mice. Empty circles: no maternal separation. Filled circles: maternally separated. (+) Mean; (Line) Median. The assessment of the main effects (L-Dopa administration and MS) was performed with Two-Way ANOVA with Tukey’s test for multiple comparison procedures. Significance was set at p < 0.05 for all analyses. (C) Correlations between latency and immobility time in Saline, L-Dopa, SKF, and Quinpirole females and males. Color intensity indicates the degree of correlation (red indicates a positive correlation and blue a negative correlation). The heatmap shows Spearman correlations and p-values. Dark borders indicate p ≤ 0.05. (D) UMAP visualization of standardized behavioral features, analyzed separately in females and males. Colors represent the four k-means clusters, whereas symbol shapes indicate treatment groups. (E) Individual heatmaps showing standardized behavioral values within each cluster. Rows represent individual animals, columns represent TST variables, and colors indicate z-scores, with red and blue denoting values above and below the mean, respectively. Side annotations identify treatment groups, and the adjacent plots summarize the average behavioral profile of each cluster.

We did not observe any significant differences in immobility time and climbing time between prepubertal males (Fig. 4H and 4J). However, we observed a main effect of maternal separation on latency, with a 30% decrease in latency in mice that underwent maternal separation (F (2,83) = 7.498, p = 0.008; Power = 0.710). We also observed a decrease of 41% in latency immobility in MS+L-Dopa males compared to L-Dopa (F (2,68) = 0,616, Two Way Anova, Tukey, p = 0.020; Power = 0,050) (Fig. 4A). In addition, there was a main effect in time and the number of immobility episodes, with mice that underwent maternal separation showing an increase of 38% in time of immobility (F (2,83) = 4.238, p = 0.043; Power = 0.426), and an increase of 56% in the number of episodes (F (2,83) = 15.707, p < 0.001; Power = 0.966). Also, there was an increase of 64% in the number of immobility episodes in MS+Saline compared to Saline (F (2,68) = 2.729, p = 0.022; Power = 0.351) and of 105% in MS+L-Dopa compared to L-Dopa (F (2,68) = 2.729, p < 0.001; Power = 0.351) (Fig. 4G). L-Dopa males showed a decrease of 64% (F (2,68) = 3.826, p < 0.001; Power = 0.537) and 48% (F (2,68) = 2.729, p = 0.048; Power = 0.351) in the number of climbing episodes compared to Naïve and Saline males respectively (Fig. 4I). Interestingly, MS+L-Dopa males had an increase of 120% in the number of climbing episodes compared to L-Dopa males (F (2,68) = 2.729, p = 0.002; Power = 0.351) (Fig. 4J).

We calculated pairwise Spearman correlation coefficients to assess the relationships among FST variables (Fig. 4K,L). All female groups, but MS+Saline, showed a correlation between climbing time and climbing frequency: Naïve (rs = 0.76, p = 0.004), Saline (rs = 0.81, p = 0.047), L-Dopa (rs = 0.90, p = 0.004), MS (rs = 0.98, p < 0.001), and MS+L-Dopa (rs = 0.84, p < 0.001) (Fig. 3K). L-Dopa females also showed a negative correlation between climbing and immobility time (rs = -0.88, p = 0.004), and MS females showed a correlation between Immobility time and immobility frequency (rs = 0.69, p = 0.048) (Fig. 4K). In males, Naïve showed negative correlations between latency and immobility frequency (rs = -0.82, p = 0.011) and immobility time (rs = -0.72, p = 0.043), and positive correlations between climbing time and climbing frequency (rs = 0.81, p = 0.011). Saline males presented negative correlations between latency and immobility time (rs = -0.66, p = 0.040), between climbing frequency and immobility frequency (rs = -0.88, p = 0.001) and immobility time (rs = -0.82, p = 0.004), and between climbing time and immobility frequency (rs = -0.80, p = 0.005) and immobility time (rs = -0.86, p = 0.002). They also presented a positive correlation between immobility time and immobility frequency (rs = 0.89, p < 0.001), and between climbing time and climbing frequency (rs = 0.90, p = 0.004). L-Dopa males showed negative correlations between latency and immobility time (rs = -0.80, p = 0.010) and between latency and immobility frequency (rs = -0.76, p = 0.011). They also showed a positive correlation between immobility time and immobility frequency (rs = 0.76, p = 0.011), and between climbing time and climbing frequency (rs = 0.81, p = 0.010). MS males presented only a correlation between climbing time and frequency (rs = 0.87, p < 0.001). Finally, MS+Saline presented a negative correlation between latency and immobility time (rs = -0.71, p = 0.046), and a positive correlation between climbing frequency and time (rs = 0.87, p = 0.002) (Fig. 4L).

UMAP visualization revealed six partially separated behavioral profiles, with treatment groups distributed across clusters (Fig. 3M,N). Individual heatmaps showed that no behavioral profile was exclusive to a group. However, females from the Naïve group were not represented in cluster 1 (higher latency and lower number and time of immobility). Interestingly, L-Dopa females were mainly present in clusters 1 and 4 (lower climbing frequency and climbing time). MS females were present in all clusters, but MS+L-Dopa were mainly present in clusters 1 and 3 (higher climbing frequency and climbing time) (Fig. 4M,N). In males, Naïve mice were absent in cluster 3 (higher immobility frequency and time), and Saline mice were present only in cluster 1 (lower climbing frequency and time) and 2 (lower immobility frequency and time). L-Dopa males were mainly present in cluster 1 and were absent in clusters 5 (higher frequency and climbing time) and 6 (higher latency and lower in the remaining parameters). MS males were mainly present in cluster 4 (intermediate in all parameters), and MS+Saline were absent in cluster 1. In addition, MS+L-Dopa males were mainly present in clusters 3 and 4 (Fig. 4O,P).

### 3.4 Early life stress, but not L-Dopa, decreased compulsive-like behavior in marble-burying test

Both dopamine signaling and maternal separation stress modulate marble-burying behavior (Jarrar et al., 2022; Jimenez-Gomez et al., 2011), which can be interpreted as compulsive-like or repetitive digging behavior (Deacon, 2006; Dixit et al., 2020). In the marble-burying test, the effects differed between females and males (Fig. 5A–B). Females that underwent maternal separation showed a main effect on the number of buried marbles, with a 21% decrease (F (2,91) = 6.627, p = 0.012; Power = 0.651). In females, the MS+Saline group showed a lower number of buried marbles compared with the Saline females (F (2,91) = 0.314, p = 0.042; Power = 0.05) (Fig. 5A). We also observed a main effect of maternal separation on the number of marbles buried in males, with a 33% decrease in marbles buried (F (2,74) = 6.370, p = 0.014; Power = 0.628). In males, MS mice buried fewer marbles than Naïve mice (F (2,74) = 0.550, p = 0.030; Power = 0.050). No other significant differences were observed among groups (Fig. 5B).

**Figure 5:**
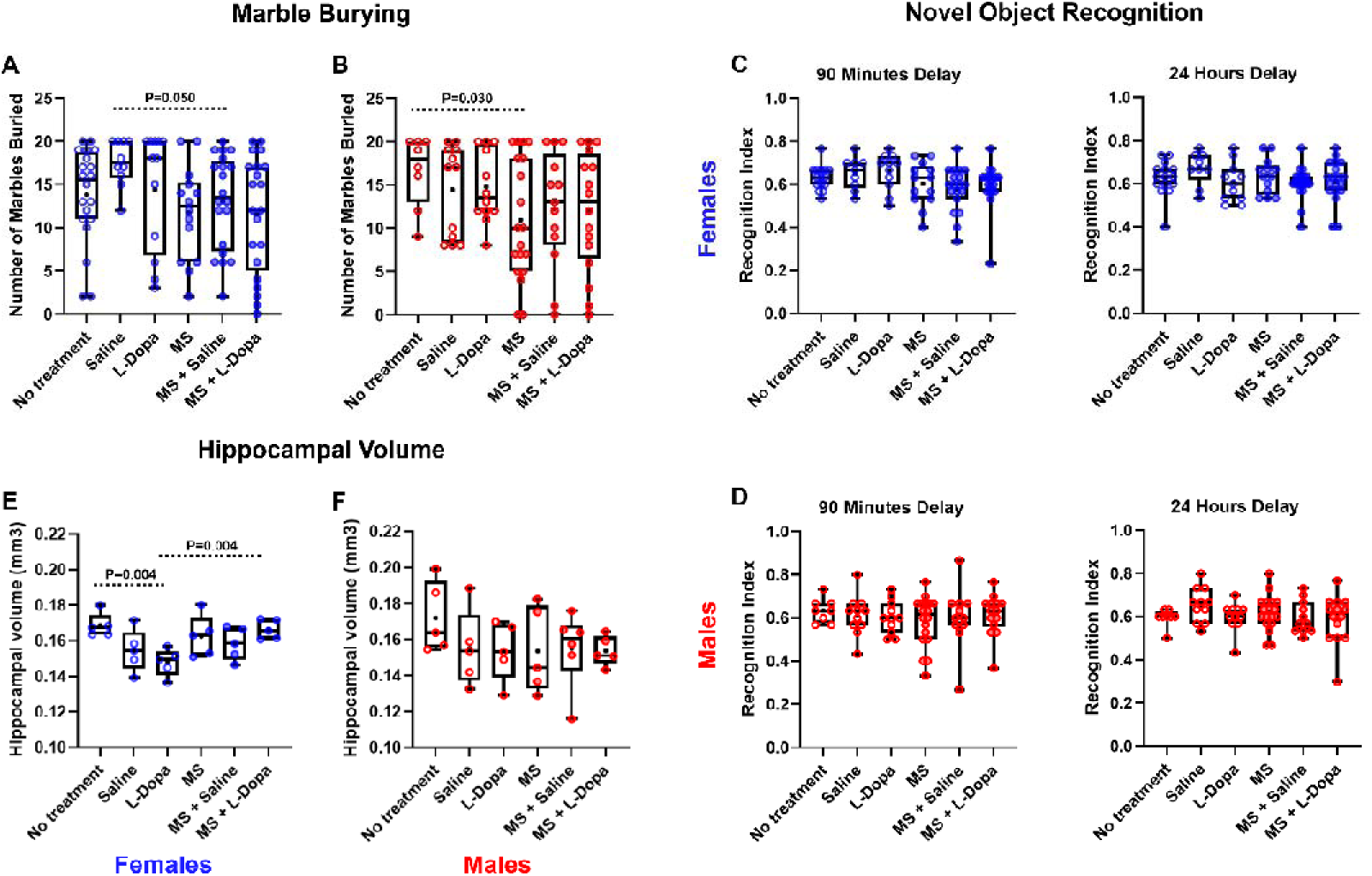
Effects of maternal separation stress and early increase in dopamine in marble burying, novel object recognition and hippocampal volume. (A,B) Compulsive-like behavior of prepubertal C57/BL6 mice in the Marble Burying Test. (A) MS+Saline females showed a decrease in the number of buried marbles compared to Saline prepubertal females. (B) MS males showed fewer buried marbles than naïve prepubescent males. (C,D) Memory of prepubertal C57/BL6 mice in the Novel Object Recognition test. (C,D) Prepubertal females and males from all experimental groups showed similar object recognition index. (E,F) Hippocampal volume was quantified using structural MRI. (E) L-Dopa females showed decreased hippocampal volume, while MS+L-Dopa showed an increase compared with L-Dopa. (F) No significant effects of treatment or sex were observed in prepubescent males. MS: Maternal Separation. Blue circles: female mice. Red circles: male mice. Empty circles: no maternal separation. Filled circles: maternally separated. (+) Mean; (Line) Median. The main effects (L-Dopa administration and MS) were assessed with Two-Way ANOVA with Tukey’s test for multiple comparison procedures. Significance was set at p < 0.05 for all analyses.

### 3.5 Early life stress and L-Dopa did not affect memory in the novel object recognition test

It is known that dopamine signaling and stress by maternal separation are involved in recognition memory (Fleury et al., 2024; Rocha et al., 2021; Titulaer et al., 2021). Thus, we assessed recognition memory using the NORT test after 90-minute and 24-hour delays (Fig. 5C–D). Regarding females, we observed a main effect in the 90-minute test in maternal separation, with females that underwent maternal separation showing a 9% decrease in memory (F (2,80) = 7.105, p = 0.009; Power = 0.685). However, we did not observe differences between female groups (Fig. 5C). We also found no differences in the 24-hour memory test (Fig. 5C). In addition, we detected no significant differences in the recognition index among males, indicating that neither maternal separation nor L-Dopa treatment significantly altered NORT in prepubertal mice (Fig. 5D).

### 3.6 Increased levels of dopamine in early development decreased female hippocampal volume

Maternal separation stress is associated with reduced hippocampal volume, and dopamine is involved in hippocampal neurogenesis (Herpfer et al., 2012; Kasahara et al., 2022). Here, hippocampal volume was affected in a sex-dependent manner (Fig. 5E–F). In females, L-Dopa mice exhibited a reduced hippocampal volume compared with Naïve (F (2,24) = 4.629, p = 0.004; Power = 0.613). However, hippocampal volume was higher in the MS+L-Dopa group than in L-Dopa females (F (2,24) = 4.629, p = 0.004; Power = 0.613). In contrast, hippocampal volume did not differ significantly among the male groups (Fig. 5F).

### 3.7 Multivariate analysis reveals sex-specific behavioral profiles across experimental conditions

To determine whether the experimental manipulations produced broader behavioral phenotypes not evident from analyses of individual variables, we performed multivariate analyses separately for females and males using standardized behavioral measures from the OFT, EPM, FST, MB, and NORT (Fig. 6).

**Figure 6:**
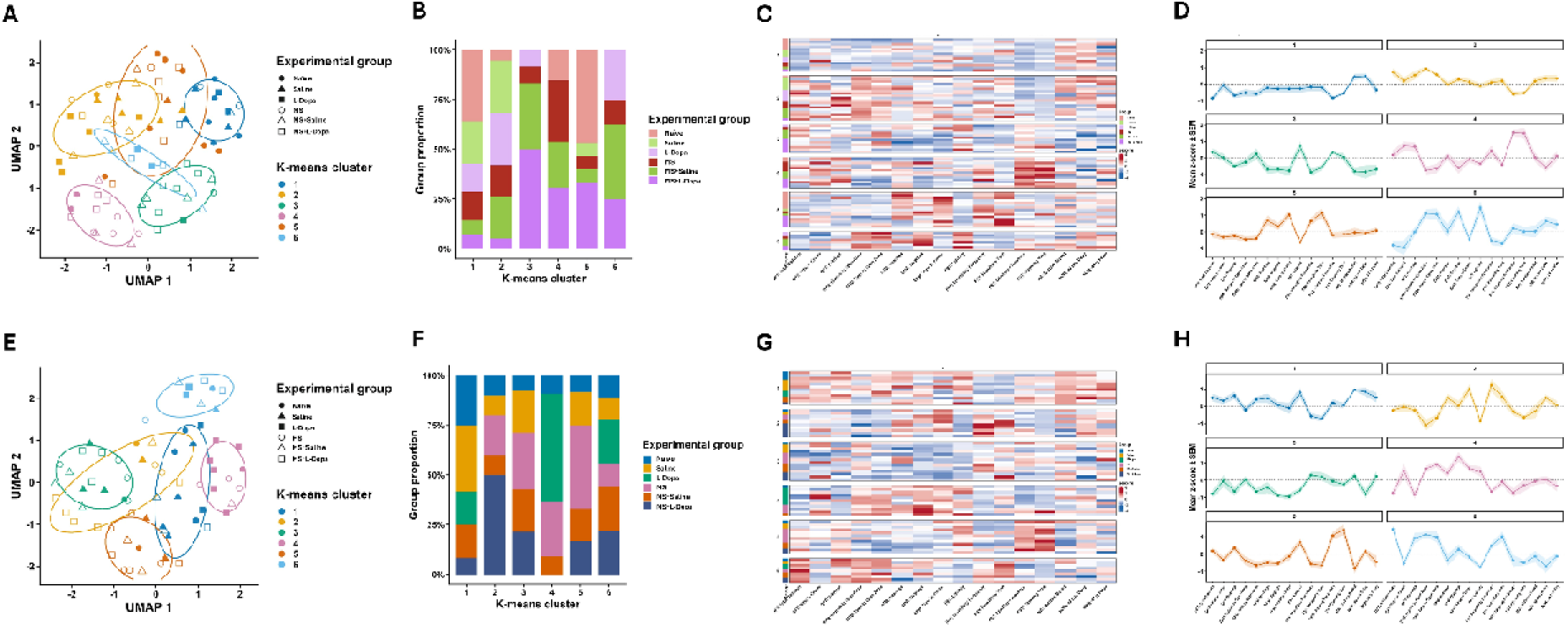
Multivariate behavioral profiles of female and male mice across experimental conditions. Multivariate analyses were performed separately for females (A–D) and males (E–H) using standardized behavioral variables from the open-field test (OFT), elevated plus maze (EPM), forced swim test (FST), marble-burying test (MB), and novel-object recognition test (NORT). (A, E) UMAP projections showing the distribution of individual females (A) and males (E). Colors indicate the six K-means clusters (k = 6), and symbols indicate experimental groups; ellipses delineate each cluster’s distribution. (B, F) Proportional composition of each K-means cluster according to experimental group in females (B) and males (F). (C, G) Heatmaps of standardized behavioral variables (z-scores) for individual females (C) and males (G), organized according to K-means cluster. Red and blue indicate values above and below the overall mean, respectively. (D, H) Mean behavioral profiles of each cluster in females (D) and males (H), expressed as mean z-score ± SEM across the behavioral variables; the dotted horizontal line indicates the overall mean (z = 0).

UMAP visualization revealed six partially separated behavioral profiles, with treatment groups distributed across clusters (Fig. 6A and E). Individual heatmaps showed that no behavioral profile was exclusive to a group. UMAP dimensionality reduction followed by K-means clustering in females (Fig. 6A). Experimental groups were distributed across multiple clusters rather than forming treatment-specific clusters, indicating substantial interindividual behavioral heterogeneity (Fig. 6B). Nevertheless, the relative representation of the experimental groups differed among clusters. Cluster 1 contained predominantly non-maternally separated animals, particularly the Naïve group, whereas Cluster 2 contained a relatively high proportion of Saline and L-Dopa females. In contrast, Clusters 3 and 6 were strongly enriched in maternally separated animals, particularly MS+L-Dopa and MS+Saline mice. Cluster 4 was also predominantly composed of maternally separated animals, whereas Cluster 5 contained a high proportion of Naïve and MS+L-Dopa animals (Fig. 6B). Interestingly, Clusters 3 and 6 contained no Naïve females, and Clusters 4 and 5 contained no L-Dopa females (Fig. 6B).

The heatmap of standardized behavioral measures showed distinct multivariate profiles among the female clusters (Fig. 6C), further illustrated by the cluster-wise mean z-scores (Fig. 6D). Cluster 1 was characterized predominantly by below-average OFT and EPM measures, along with relatively higher FST climbing measures. Cluster 2 displayed generally elevated exploratory measures in the OFT and EPM, including open-arm exploration, together with comparatively reduced FST climbing. Cluster 3 showed reduced values for several EPM measures and marble burying, accompanied by increased FST latency. Cluster 4 was distinguished by markedly increased FST climbing frequency and climbing time, together with relatively reduced open-arm exploration. Conversely, Cluster 5 was characterized primarily by higher FST immobility measures, whereas Cluster 6 showed increased EPM open-arm exploration and head dipping, increased FST latency, and comparatively low immobility measures. Thus, the female clusters separated females mainly by combinations of exploratory and stress-coping behaviors rather than by experimental treatment alone.

Males showed a similarly heterogeneous but distinct organization (Fig. 6E–H). As in females, no cluster corresponded exclusively to a single experimental condition (Fig. 6F). However, experimental treatments were differentially represented among the clusters. Cluster 1 was predominantly composed of non-maternally separated animals, whereas Clusters 2, 3, and 5 contained a greater proportion of maternally separated mice. Cluster 4 was particularly enriched in L-Dopa animals, while Cluster 6 showed a more heterogeneous distribution of all experimental conditions.

The standardized behavioral profiles also differed markedly across male clusters (Fig. 6G,H). Cluster 1 exhibited generally above-average OFT exploratory measures, reduced FST immobility, and relatively high marble-burying and NORT scores. Cluster 2 showed a mixed phenotype characterized by low OFT rearing but increased head dipping, FST latency, and some immobility-related measures. Cluster 3 displayed generally reduced exploratory measures and comparatively low 90-min NORT performance. Cluster 4 was characterized by pronounced EPM open-arm exploration and head dipping, together with relatively low FST immobility and climbing measures. Cluster 5 displayed reduced EPM open-arm exploration but markedly elevated FST climbing frequency and climbing time, whereas Cluster 6 combined increased OFT locomotion and EPM exploration with greater FST immobility time and reduced 90-min object-recognition performance.

Overall, the multivariate analysis organized behavioral phenotypes into distinct combinations of exploratory, anxiety-like, stress-coping, repetitive-digging, and recognition-memory measures. Experimental groups were distributed across several clusters, but their relative enrichment differed between clusters and between sexes.

## Discussion

Several neuropsychiatric disorders share both dopamine abnormalities and early life stress hypotheses (Ijomone et al., 2024). Because stress and dopaminergic signaling interact bidirectionally (Anderson et al., 2026; Brake et al., 2004; Pruessner, 2004), we investigated whether L-Dopa administration during maternal separation affects behavioral development at prepuberty. Here, we showed that exposure to a transient pharmacological increase in dopaminergic signaling and maternal separation during early postnatal development produce persistent, domain-specific, and sex-dependent behavioral effects detectable during prepuberty. Locomotor activity was preserved, recognition memory remained relatively intact, and multivariate analyses showed overlap among experimental groups rather than treatment-specific behavioral clusters. Instead, the effects were selective to exploratory behavior, elevated plus maze responses, coping with an acute inescapable stressor, marble-burying behavior, and hippocampal volume. L-Dopa treatment from PD1 to PD14 increased rearing in OFT in females and modified EPM and FST behaviors in both sexes, whereas maternal separation altered several of these effects when combined with L-Dopa. Maternal separation also reduced marble burying in specific comparisons, without impairing novel-object recognition. Finally, we detected reduced hippocampal volume specifically in L-Dopa females, while hippocampal volume was preserved in males and was higher in MS+L-Dopa females than in females exposed to L-Dopa alone. Together, these findings indicate that early increases in dopamine availability and early stress alter the developmental organization of specific behavioral domains. The extensive overlap observed in the multivariate analyses further suggests that these developmental manipulations shift the probability of particular behavioral profiles rather than producing discrete treatment-defined phenotypes.

### Dopamine increase during infancy leads to exploratory and coping behavior in prepubertal females, and decreases anxiety-like and coping behavior in prepubertal males

We previously showed that perinatal L-Dopa administration increases dopamine levels within the brain of pups (de Matos et al., 2018). Here, L-Dopa treatment alone produced different effects in prepubertal females and males. In females, an early increase in dopamine levels augmented rearing in the OFT without affecting total distance traveled or time in the center, suggesting a selective alteration in vertical exploratory activity rather than generalized hyperlocomotion. Since there were no correlation between total distance traveled and rearing frequency in females, we interpret that the L-Dopa increase in female rearing is not simply secondary to generalized hyperlocomotion. Instead, developmental L-Dopa appears to have selectively affected the vertical/exploratory component of OFT behavior, which could reflect environmental investigation, novelty engagement, or behavioral information gathering (Borta and Schwarting, 2005; Lever et al., 2006; Sturman et al., 2018). These results are similar to our previous study, in which 5 postnatal days of L-Dopa administration also increased exploratory behavior only in females (de Matos et al., 2018). In the EPM, L-Dopa females showed more open-arm entries and spent more time in the open arms than Naïve females. This could be interpreted as an anxiolytic-like behavioral profile, but the similar increase observed in Saline females indicates that part of this phenotype may be attributable to repeated neonatal injection/handling rather than specifically to L-Dopa. In contrast, both parameters were positively correlated in L-Dopa but not in Saline, indicating that L-Dopa females that entered the open arms more often also tended to remain there longer. Thus, the L-Dopa phenotype is behaviorally more coherent than the Saline phenotype. Interestingly, we did not see any EPM changes in 5 postnatal-day L-Dopa administration (de Matos et al., 2018). In the FST, L-Dopa females displayed fewer immobility episodes and less total immobility than Saline females. These alterations can be interpreted as resilience to despair/depressive-like behavior, or a shift away from passive coping with an acute inescapable stressor (Castagné et al., 2009; Commons et al., 2017; Slattery and Cryan, 2012). In addition, climbing was strongly and negatively associated with immobility time, meaning prepubertal females that expressed more active coping through climbing tended to spend less time immobile in passive coping. Again, we did not see FST changes when L-Dopa was administered only in the first 5 postnatal days (de Matos et al., 2018).

In males, L-Dopa did not alter OFT or rearing but increased time in the open arms and head dipping relative to Saline, suggesting anxiolytic-like behavior or enhanced engagement with the exposed/aversive portions of the maze. Conversely, L-Dopa males displayed fewer climbing episodes during FST without corresponding changes in total immobility or climbing time. Because climbing represents active, escape-directed coping, this result suggests a decrease in the pattern or initiation of active coping (Detke et al., 1995; Slattery and Cryan, 2012). Thus, early L-Dopa exposure appears to reorganize behavioral responses differently in females and males. This developmental specificity is consistent with previous work from our group showing that early postnatal L-Dopa exposure produces behavioral effects that depend on both sex and age at assessment (de Matos et al., 2020, 2018).

### Prepubertal MS females showed anxiolytic-like active coping behavior, while maternal separation induced anxious-like and passive coping behavior in prepubertal males

In the OFT, MS did not alter locomotor or exploratory activity, whereas the EPM revealed divergent responses between sexes: MS females spent more time in the open arms, while in males, as a main factor, maternal separation was associated with reduced open-arm time. This divergence extended to coping behavior in the FST. In females, as a main factor, MS increased both the frequency and duration of climbing, suggesting greater engagement in active escape-directed coping, whereas in males, as a main factor, MS reduced latency to immobility and increased immobility frequency and duration, indicating earlier and greater recruitment of passive coping. These sex-dependent outcomes are consistent with evidence that the behavioral consequences of maternal separation vary substantially by sex and developmental stage. Indeed, studies in adolescent C57BL/6 mice have reported stronger passive-coping effects of early-life stress in males and different or weaker effects in females (He et al., 2020). Importantly, as a main factor, maternal separation also reduced marble-burying behavior in both sexes, which is more conservatively interpreted as reduced repetitive digging/burying or compulsive-like behavior rather than reduced anxiety (Angoa-Pérez et al., 2013). Recognition memory was preserved, except for a main effect of maternal separation on the 90-min recognition index in females, with no significant pairwise differences, and MS alone did not significantly alter hippocampal volume. This selective pattern is consistent with systematic and meta-analytic evidence showing substantial heterogeneity in mouse maternal separation phenotypes and relatively weak or inconsistent effects in conventional OFT and EPM measures, emphasizing that the consequences of maternal separation depend on protocol, sex, species, and age at assessment (Stupart et al., 2023; Tractenberg et al., 2016; Wang et al., 2020).

### MS+L-Dopa attenuates dopamine increase and early-life stress effects on OFT and EPM, but has a putative additive effect on despair/depressive-like behavior in males

As said before, dopaminergic abnormalities and early-life stress are both implicated in the pathophysiology of several neuropsychiatric disorders. In this study, we pharmacologically increased dopamine levels during stress by maternal separation to evaluate their interaction during postnatal development. We designed two hypotheses: (1) dopamine increase and early-life stress would have additive effects in prepubertal behavior; (2) dopamine increase and early-life stress would attenuate each other’s effects on behavioral development. We observed that, in females, maternal separation abolished the L-Dopa-associated increase in OFT rearing and reduced time spent in the open arms relative to L-Dopa females in the EPM. Interestingly, in MS+L-Dopa females, center time correlated positively with head dippings. Since the central platform is an important decision/transition zone in the EPM, and head dipping is often interpreted as active risk assessment, one possible explanation is that MS+L-Dopa females spending more time in the decision/transition region also engaged more strongly in active environmental/risk assessment. In the FST, MS+L-Dopa females showed more immobility episodes and longer immobility duration than L-Dopa females, while also showing more climbing episodes. Together, these results may reflect despair/depressive-like behavior or altered organization of coping behavior, with more frequent transitions between active and passive responses. These results showed that, in females, MS+L-Dopa attenuated the L-Dopa effects. Furthermore, the gross structural findings are also noteworthy because MS+L-Dopa females had larger hippocampal volumes than L-Dopa females, effectively eliminating the reduction detected after L-Dopa alone.

In males, a similar interaction was evident. MS+L-Dopa males spent less time in the open arms and showed fewer head dips than L-Dopa males in EPM, had a shorter latency to immobility and more immobility episodes, but also showed more climbing episodes than L-Dopa males. These results indicate that maternal separation also attenuated the behavioral consequences of developmental L-Dopa exposure in males. However, one interesting result is highlighted: MS+L-Dopa males showed shorter latency to immobility and more immobility episodes in FST. These behavioral phenotypes are not altered in L-Dopa males, which suggests putative additive effects of L-Dopa and maternal separation on despair/depressive-like behavior in prepubescent males.

### Distinct effects in females and males

In the present study, females were more sensitive to L-Dopa-induced changes in exploratory behavior and were the only sex in which hippocampal volume was altered, whereas males exhibited a prominent L-Dopa-associated change in climbing frequency. Likewise, maternal separation interacted differently with behavioral outcomes according to sex. Total distance traveled and rearing frequency were positively correlated in several male groups, whereas OFT showed no significant correlations in females. This pattern suggests that horizontal locomotion and vertical exploration are behaviorally coupled in males, whereas in females these dimensions appear more independent. Another interesting result was noticed in the EPM. In females (except Saline), time in open arms and the number of entries in the open arms were positively associated, but not in males (except MS+L-Dopa). Thus, we observed this evidence of internal behavioral coherence only in prepubertal females. Such sex dependence is consistent with studies showing that early-life stress can produce different developmental trajectories in female and male mice. Importantly, these differences may emerge at distinct ages rather than representing stable sex differences throughout life. Goodwill et al., for example, found sex-selective consequences of early-life stress that developed progressively from adolescence to adulthood (Goodwill et al., 2019). The clear divergence between females and males reinforces the importance of considering sex as a biological variable in studies of developmental stress and dopamine signaling. Dopaminergic and corticolimbic systems do not follow identical maturational trajectories in females and males, and sex differences in hippocampal, amygdala, and prefrontal development emerge well before adulthood. These differences are influenced not only by gonadal hormones at the time of behavioral testing but also by perinatal hormonal organization, sex-chromosome effects, developmental differences in receptor expression and synaptic maturation, and sex-specific regulation of the HPA axis (Premachandran et al., 2020).

### Prepubertal behavior

The age at which the mice were tested is central to interpreting these findings. Behavioral testing between PD28 and PD32 evaluates the consequences of PD1–14 manipulations before complete maturation of the dopaminergic, corticolimbic, and stress-regulatory systems. The period immediately after weaning is characterized by extensive remodeling of reward, motivational, prefrontal, hippocampal, and stress-responsive circuits, while dopaminergic innervation and dopamine-dependent regulation of cortical neurons continue to mature through adolescence. For example, dopamine modulation of behavior or prefrontal interneurons changes across adolescence, and mesocortical dopamine projections undergo substantial developmental refinement during this period (Moraes et al., 2024; Peters and Naneix, 2022). It is worth noting, however, that PD28–32 should not necessarily be considered a uniformly prepubertal interval in C57BL/6 mice. External markers of pubertal onset can appear around PD27–29, although complete reproductive maturation occurs later and varies substantially among individuals (Hou et al., 2022). Thus, the animals in the present study are more precisely situated in a late juvenile/early peripubertal transition, a developmental stage in which hormonal, neural, and behavioral systems change rapidly. This may help explain why the observed effects differ from those previously identified in older animals after similar neonatal dopaminergic manipulation (de Matos et al., 2020). It also raises the possibility that some phenotypes observed here represent transient developmental alterations that could disappear, reverse, or intensify during adolescence and adulthood. Longitudinal or age-comparative studies will therefore be important to determine whether the increased female rearing, altered coping strategies, and reduced female hippocampal volume represent temporary developmental delays, accelerated maturation, or persistent alterations in behavioral trajectories. The study is particularly informative because most maternal-separation studies evaluate animals only after adolescence or in adulthood, potentially missing phenotypes specific to this transitional developmental period.

### Dopaminergic imbalance and early-life stress “two-hit model”

These findings can also be considered within neurodevelopmental models that propose interactions between dopamine dysregulation and adverse developmental experiences. Dopamine is not only a mature neurotransmitter but also participates in neurodevelopmental processes, including progenitor proliferation, neuronal migration, GABAergic differentiation, maturation of striatal projection neurons, and establishment of cortical circuitry (Crandall et al., 2007; de Souza Lima et al., 2020; Souza et al., 2024, 2011). At the same time, early-life adversity can modify the HPA axis, corticolimbic development, and mesocorticolimbic dopamine function, potentially altering later responses to stress and motivational stimuli. Maternal rearing conditions have been shown to influence mesocorticolimbic dopamine responses, and human studies also support interactions between early maternal care, stress, and dopamine release. These observations provide a biological intersection between the dopamine hypothesis of neuropsychiatric dysfunction and developmental stress hypotheses. The present results are also relevant to the conceptual framework of the “two-hit” model, in which an early perturbation alters brain development and creates a substrate that responds to a second perturbation differently from an otherwise normally developing brain (Maynard et al., 2001). However, the current experiment should not be described as a strict test of the classical two-hit hypothesis, because L-Dopa administration and maternal separation occurred during the same PD1–14 developmental interval, rather than as temporally separated first and second hits. It therefore more directly represents a dual-perturbation or convergent developmental-hit model. Nevertheless, the results provide an important mechanistic insight relevant to two-hit theories: developmental insults need not combine additively. In fact, MS frequently attenuated, reversed, or qualitatively changed L-Dopa-associated phenotypes, including exploratory behavior, EPM behavior, FST coping patterns and female hippocampal volume. This supports contemporary extensions of the two-hit concept in which interacting biological and environmental exposures alter developmental trajectories through nonlinear processes rather than simply accumulating damage (Davis et al., 2016). Accordingly, dopamine dysregulation may modify the neural substrate on which stress acts, while early stress may simultaneously reshape dopamine-dependent circuit maturation. Their interaction could therefore determine how behavioral phenotypes emerge rather than simply how severe they become.

### Limitations

Several limitations should be considered. First, we used L-Dopa plus benserazide to increase central dopamine availability, but we did not directly measure dopamine concentrations, turnover, receptor expression, or downstream signaling. Thus, we cannot yet link the persistence of the behavioral effects to a specific dopaminergic mechanism, receptor subtype, or circuit. Moreover, L-Dopa can ultimately influence other catecholaminergic pathways, preventing attribution of all effects exclusively to dopamine. Second, repeated intraperitoneal injections constitute a developmental manipulation themselves and can affect behavior (Moraes et al., 2025). Indeed, several differences between Saline and Naïve animals in the EPM and FST demonstrate that injection and/or repeated handling was not behaviorally neutral and emphasize the importance of retaining both control groups when interpreting L-Dopa effects. Third, maternal separation was performed while littermates remained together and were maintained under warm conditions, potentially reducing the intensity of the stressor relative to protocols involving individual isolation; variation in duration, temperature, dam behavior, litter composition and post-separation maternal care is known to contribute substantially to the heterogeneity of maternal-separation outcomes (Tractenberg et al., 2016). Fourth, endocrine markers of stress such as corticosterone, as well as maternal behavior following reunion, were not measured, limiting our ability to establish the physiological magnitude of the early-life stress exposure. Finally, pubertal status was not directly determined. Because PD28–32 overlaps with the onset of puberty in C57BL/6 mice, individual differences in vaginal opening, estrous development or preputial separation may have contributed to behavioral variability. Replication across developmental ages, together with neurochemical, endocrine and circuit-level analyses, will be necessary to establish the mechanisms underlying the behavioral interactions identified here.

### Conclusion

In conclusion, our findings indicate that transient modification of dopamine availability and maternal environment during the first two postnatal weeks can alter behavioral development in prepuberty, but their effects depend strongly on sex, behavioral domain, and the interaction between the two developmental experiences. Early L-Dopa exposure selectively increased exploratory behavior in females, altered EPM behavior and coping strategies in both sexes, and reduced hippocampal volume specifically in females, while preserving recognition memory. Maternal separation reshaped many of these L-Dopa-associated phenotypes, frequently opposing rather than potentiating them, and restored hippocampal volume in L-Dopa-treated females. Thus, the consequences of simultaneous dopaminergic and stress-related perturbations cannot be predicted from either exposure alone. Rather than supporting a simple cumulative “two-hit” mechanism, the data suggest that dopamine signaling and early-life stress interact dynamically during sensitive periods of brain maturation to redirect subsequent behavioral trajectories. Differences between females and males, and the emergence of these effects around the juvenile-to-pubertal transition, further emphasize that developmental stage and sex fundamentally determine the consequences of early environmental and neurochemical perturbations. These findings support a model in which vulnerability to later neuropsychiatric phenotypes arises not from a single developmental insult, but from interactions among neurochemical state, environmental experience, sex, and developmental timing.

## Acknowledgments

To the young scientists at the NeuroDEv laboratory, thank you for your daily support. To our colleagues, we appreciate our scientific discussions and the assistance you provided in times of urgency. And a special acknowledgment to the often “invisible” yet indispensable members of our university community - the technicians, cleaning personnel, security team, restaurant staff, and administration professionals - your contributions are the bedrock of our research endeavor.

